# Highly persistent antibody levels but limited population immunity in gannets after HPAI outbreak

**DOI:** 10.64898/2026.08.25.745523

**Authors:** Thierry Boulinier, Mathilde Lejeune, Pascale Massin, Eric Niqueux, Armel Deniau, Raphaëlle Woerlé, Alice Bernard, Aurore Ponchon, Tristan Martin, Jérôme Fort, Béatrice Grasland, David Grémillet, Pascal Provost, Jérémy Tornos

## Abstract

The recent large-scale circulation of High Pathogenicity Avian Influenza (HP AI) viruses H5Nx of clade 2.3.4.4b has been responsible for massive die-offs in wild species, notably in long-lived seabirds, with unknown implications for the immunity of surviving individuals. In the North Atlantic, northern gannet colonies were heavily affected in 2022, with more than 40% mortality observed among breeding adults and some surviving individuals developing dark irises. Using samples collected in 2023 and 2024 on Rouzic colony (France), we report persistent individual anti-AI antibody levels and seroneutralisation titres, with most of the immune individuals showing dark irises. A modelling approach further stressed the importance of long-lasting immunity in such species by showing that the proportion of individuals which kept their immunity between years strongly limited decreases in population size in case of repeated outbreaks. Overall, our results highlight the existence and importance of long-lasting immunity in long-lived species for population persistence.

## Introduction

Determining which individuals survive deadly epizooties and why is critical for predicting the effects of pathogens on host populations (Keeling & Rohani 2007). This knowledge is particularly important for the conservation of wild populations in a context of increasing emergence of pathogens (Daszak et al. 2000; Foufopoulos et al. 2022; Rustadler 2024). One way to obtain this information is to quantify the presence of antibodies against a pathogen among individuals over time after epizooties (Pepin et al. 2017). Antibodies are molecules of the vertebrate immune system that are produced within a few weeks of exposure to an antigen. Their persistence in the blood is highly variable, depending on the host, pathogen, and time since exposure (Tizard 2013). Relatively little is known about the temporal persistence of antibody levels against disease agents in wild species (Gilbert et al. 2013; Gamble et al. 2020; Wight et al. 2024), but long-lived species have been suggested to have evolved long-lived acquired immunity due to their life history (Garnier et al. 2013). The detection of antibodies in blood samples of surviving individuals can reveal the proportion of individuals that have been exposed to a pathogen (Hens et al. 2012). An association between antibody level and actual protection against specific pathogens is nevertheless not always detected (Caliendo et al. 2022). Nevertheless, in some host-pathogen interactions, antibodies play a key role in the protection of host individuals, and their detection can inform us on the immune response that may have allowed individuals to survive, and that may protect them in case of future possible exposure (Tizard 2013).

Since 2021, epizooties of High Pathogenicity Avian Influenza (HPAI) H5Nx viruses of sublineage 2.3.4.4b have led to the death of millions of wild and domestic birds on all continents (Klaasen & Wille 2023; Gamarra-Tolledo et al. 2023; Kuiken et al. 2026). The geographic extent, the number of individuals exposed and the range of species have reached unprecedent levels (Klaasen & Wille 2023). Among wild birds, scavengers, but also colonial breeding seabirds have been heavily impacted (Falchieri et al. 2022; Banyard et al. 2022; Knief et al. 2024; Avery-Gomm et al. 2024). In particular, during spring 2022, most breeding colonies of northern gannets *Morus bassanus* (gannets, hereafter), have been seriously affected in the North Atlantic (Lane et al. 2023), with high proportions (>40%) of breeding birds dying at some colonies. Because this seabird breeds in discrete and often large aggregates where monitoring can be conducted consistently, an international effort enabled the gathering of extensive and detailed epidemiological data across most of the species’ breeding range (Lane et al. 2023; Grémillet et al. 2023; Briand et al. 2025; Matthiopoulos et al. 2026).

It has been suggested that the virus has become enzootic in wild populations at broad spatial scales (Pohlmann et al. 2022 and, despite limited knowledge about transmission processes among and within populations (Boulinier 2023; Clessin et al. 2025), a pressing question is whether long-lasting immunity of individuals could limit the impact of the disease in case of repeated local introductions of the virus. This is notably for colonial species like gannets, which are threatened in the context of global change (Jeglinski et al. 2024), breed densely, and have a seasonal use of their colonies. To date, very little information is available on the long-term persistence of antibody level against HAPI in long-lived wild species, which is also critical for interpreting serological monitoring data (Günther et al. 2024; Greco et al. 2025; Knief et al. 2026).

One striking feature of the impact of HPAI in gannets was documented early on: some individuals showed black irises, a phenotypic change potentially associated with HPAI infection (Lane et al. 2023). The physio-pathological process underlying this condition is not fully elucidated yet, but during the 2022 HPAI outbreak, one field study carried out in the largest gannet colony of Bass Rock, Scotland (Lane et al. 2023) and one carried out in the unique French colony of gannets, in Rouzic, (Grémillet et al. 2023) suggested that gannet iris phenotype could constitute a practical way to identify birds previously infected by HPAI. The occurrence of black-eyed individuals on a colony may reflect survival after a HPAI infection, but it is unclear what proportion of surviving infected individuals they represent, and whether they could be immune to reinfection (Lane et al. 2023; Petalas et al. 2025).

In this context, the present study aimed at evaluating the impact of the 2022 HPAI outbreak on the immunological profiles of individuals and their implications. To do so, we analysed blood samples from juvenile and adult breeding gannets of Rouzic Island, France collected during two consecutive breeding seasons (2023 and 2024) following the HPAI outbreak of 2022 and one year prior to the outbreak (2019). The repeated sampling of individuals allowed us to show evidence of highly persistent immunity between years. We used a model to highlight the implications of this result.

## Materials & Methods

### Study population and field sampling

Gannets were studied on Rouzic Island, Brittany, France (48°54′N, 3°26′W), where they breed between February and October, with hatching peaking during the first half of June. The main fieldwork was conducted during July-August 2023 and June 2024, one and two years respectively after the dramatic outbreak of HPAI (Grémillet et al. 2023, Scoizec et al. 2024). In addition, we took advantage of blood samples collected on 20 gannets in 2019 to explore potential signs of historical exposure to avian influenza H5Nx virus.

A total of 37 breeding adults and 10 juveniles were captured and sampled within the colony in 2023, in addition to 28 breeding adults in 2024, including 11 birds sampled in 2023. During both field sessions, birds were handled under strict biosecurity measures. Breeding adults were selected among gannets incubating a 5 weeks-old chick, while juveniles were captured on nest, ready to fledge. Bird selection focused on catching balanced numbers of gannets with normal, pale, iris *versus* with altered, darkened, iris. This led to defining three iris phenotypes: adults with pale (normal) iris on both eyes were classified as healthy-like gannets (“pale iris” group); adults with one or two full-black irises (”full-black iris” group; **Figures 1A and 1C**); adults with one or two black spotted irises (“spotted-black iris” group; **Figures 1B and 1D**). All juveniles displayed healthy-like irises and no phenotypic change was noticed during or after the outbreak. No field observation was available for the 20 adult gannets sampled in 2019 but the black-iris condition had never been documented before 2022 despite long term monitoring on the colony (Ponchon et al. 2026), so we assumed those birds displayed pale irises. **Supplementary Table 1** details the sampling size for each iris-phenotype group and sex for 2023 and 2024.

**Figure 1.**
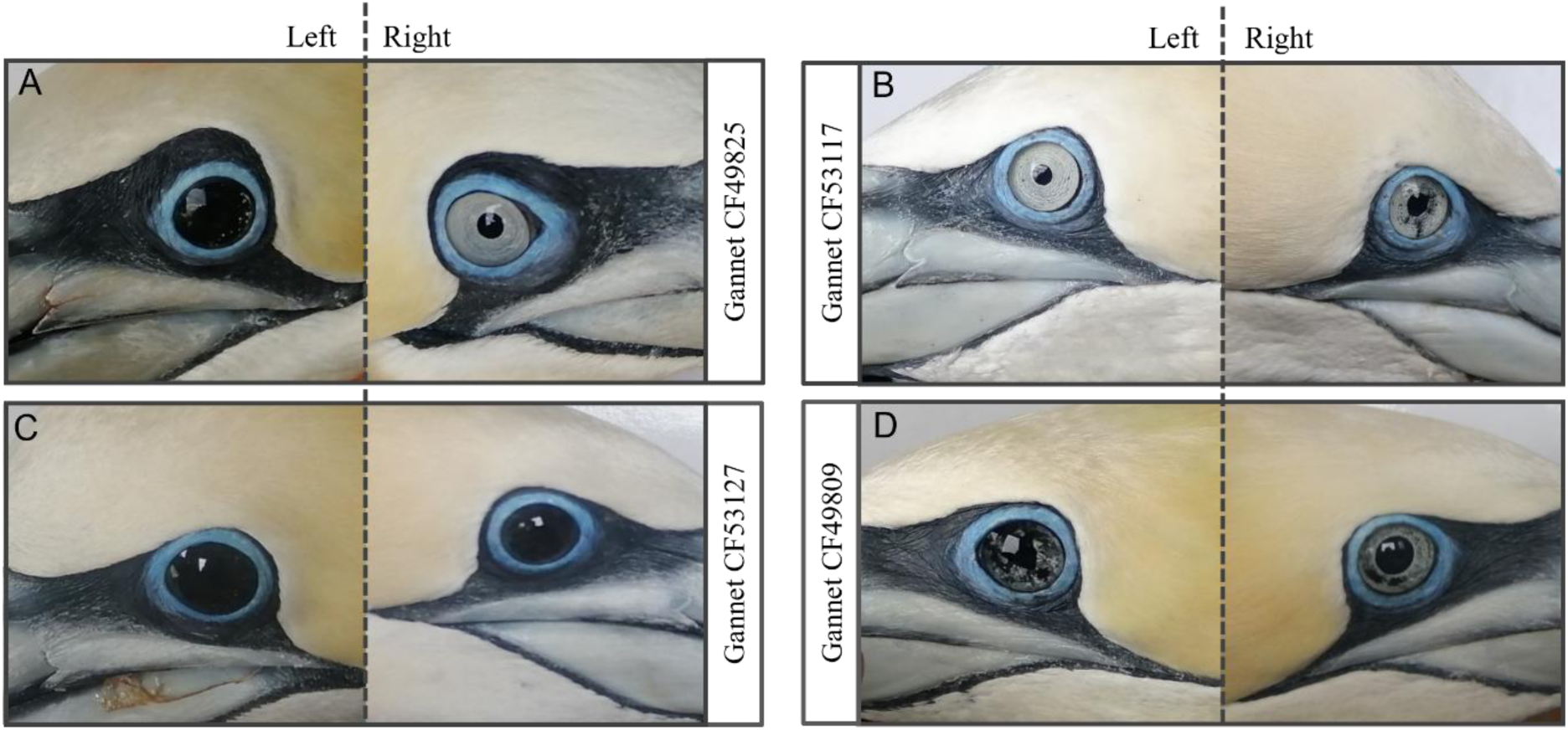
Characterisation of the different iris phenotypes observed on sampled gannets in the colony of Rouzic island, after the 2022 HPAI outbreak. Pictures refer to the left and right profiles of four different gannets as examples of the different kinds of irises we observed on the field: (**A**) gannet with a left full-black iris and a right healthy-like pale iris; (**B**) gannet with a right spotted-black iris and a left healthy-like pale iris; (**C**) gannet with two full-black irises; (**D**) gannet with two spotted-black irises. Each bird is associated with a unique alphanumeric code (e.g., CF49825) used for bird identification.

Blood samples (1 to 2 ml) were taken from the tarsal vein using a 2.5 ml syringe previously rinsed with anticoagulant (heparin). Plasma was obtained after blood centrifugation within an hour of blood collection. Plasma samples were kept cool in the field, then stored at −20°C once back at the laboratory until serological analyses. A dry cloacal swab was taken from each individual for direct HPAI virus detection. Swabs were kept cool in the field and were directly driven to the French avian influenza reference laboratory (ANSES, Ploufragan) after each field session to conduct RtPCR analyses. A few body feathers were also collected to allow molecular genetic sex determination of each individual. In addition, five adult gannets with full-black iris (unknown breeding status), found wrecked on the Brittany coast, were blood sampled in a wildlife rescue center near the colony (LPO Ile Grande rescue center) during winter 2023 for plasma analyses.

### Evaluation of the proportion of individuals with at least one full-black or spotted-black iris in the colony

Breeding success is monitored each year on a plot using an on-site remote video camera (CCTV system; Ponchon et al. 2026). In 2023 and 2024, this allowed to characterize the iris colour phenotype of both eyes of a fraction of breeding individuals over 102 nesting sites. For most of breeding sites, the iris colour phenotype of both eyes could be characterized for only one member of a breeding pair.

### Detection and quantification of anti-AIV antibody levels

Commercially-available enzyme-linked immunosorbent assay (ELISA) kits were used to detect anti-NP and anti-H5 antibodies against avian influenza viruses in plasma samples as in Lejeune et al. (2026) and as detailed in **Supplementary methods**. We used kit threshold values to discriminate seropositive from seronegative samples (NP-cELISA = 0.45; H5-cELISA = 0.50; H5-iELISA = 0.50).

### HP H5 seroneutralisation assay

Seroneutralisation assays were run in duplicate on each plasma sample to explore the capacity of specific antibodies to seroneutralise a HPAI H5 virus of clade 2.3.4.4b infection. The assays were run as in Lejeune et al. (2026) and as detailed in **Supplementary Methods**. Titres equal or above 16 (log2 = 4) were considered as seropositive.

### Statistical analyses

Statistical analyses were conducted using R (R Core Team, 2023). Observed NP and H5 antibody prevalences (P_observed_) were calculated from the ELISA’s outcomes, according to the seropositivity threshold for each ELISA assay. NP and H5 antibody true prevalences (P_true_) were estimated by considering the sensitivity (sens) and the specificity (spec) of each ELISA assay, and were calculated following: P_true_ = [P_observed_-(1-spec)]/(sens+spec–1). According to each ELISA kit documentation, the sensitivity and the specificity were both set to 0.99 for NP-cELISA, H5-cELISA and H5-iELISA. Associated 95% confidence intervals were calculated using the Clopper-Pearson estimation. Proportions of individuals with variable serological statuses were compared as a function of their iris phenotypic appearance and the sampling field season, using Two-Ways Analyses of Variance and Tukey multiple pairwise comparisons. Linear regressions and pearson correlations were used to compare the titres obtained with the different assays.

We analysed antibody levels using Gaussian linear models. Three models were used to explore potential associations between 3 dependent variables (namely the probability to be seropositive for NP, the probability to be seropositive for H5 with the competitive H5 ELISA, and the probability to be seropositive for H5 with the indirect H5 ELISA) and the iris phenotype and sex. Iris phenotype was coded either as a binary factor (pale *versus* dark = full-black + spotted-black) or as a three-level factor (pale, spotted-black, full-black), controlling for sex. Competing specifications additionally included a year effect and an iris × year interaction. Models were fitted by maximum likelihood and compared using AICc (MuMIn), reporting ΔAICc and Akaike weights. Estimated marginal means and 95% confidence intervals (CI) were obtained with *emmeans*, with Tukey-adjusted pairwise contrasts for three-level iris models. An effect of breeding stage (adult versus juveniles) was also tested. Model assumptions were checked via residual diagnostics.

### Eco-epidemiological modelling

A simple S(EI)R (Susceptible [Exposed Infected] Resistant) model was used to highlight how knowledge of the temporal persistence of immunity is important to interpret and predict changes in proportions of seropositive individuals and in population size under scenarios of re-exposure to the disease agent. Among key eco-epidemiological parameters to consider, we characterized the temporal persistence of immunity by defining OMEGA as the proportion of individuals that remained immune from one year to the next. While epidemiological data were gathered only over a time lap of one and two years after the HPAI outbreak, knowledge of demographic parameters is available over longer time periods in gannets (Gremillet et al. 2020, Lane et al. 2026).

The model allowed us to keep track in-between years of Susceptible and Resistant (immune) individuals over time. Given the high seasonality of the presence of gannets in the colony and the delayed age at first reproduction in a long-lived seabird such as the gannet, a (Leslie) matrix approach (Caswell 2006) was used to implement temporal changes in the vector of abundance of individuals in each age class (one-year old immatures, 2-year-old immatures, 3-year-old immatures, and breeding individuals). The Susceptible and Resistant compartments were tracked at each (yearly) time step at the end of the breeding season, Exposure, Infection and death from infection being assumed to occur within the breeding season. Details of the model and R code for the computations is available in the online supplement (**Supplementary information Model**).

Three key epidemiological parameters were considered: BETA, the proportion of Susceptible individuals that get infected a year of outbreak, GAMMA, the proportion of infected individuals that become Resistant (immune) if infected a year of outbreak, and OMEGA, the proportion of Resistant individuals that maintain their immunity each year and thus stay Resistant.

## Results

### Eye phenotypes in Rouzic colony

No individual with a dark-iris phenotype had been documented prior to the HPAI outbreak of 2022. In 2022, some individuals were detected with only one or two full-black irises (Figure 1A and 1C), some had one full-black iris and one iris with black spots, some had one or two irises with black spots (Figure 1B and 1D) and the rest had two pale irises.

In 2023, of the 102 monitored nests, 86 were occupied. Among the 103 individuals for which both irises were characterized, 77.7% (80/103) had two healthy-like pale irises, while 22.3% (23/103) had at least one full-or spotted-black iris - specifically, 9/23 individuals had one or two spotted/black irises and 14/23 had one or two full-black irises. In 2024, the same 102 nests were monitored, yielding similar proportions. Only one nest was unoccupied. Both irises’ patterns were recorded for 169 individuals. Among them, 76.3% (129/169) had two healthy-like pale irises, while 23.7% (40/169) had at least one full-or spotted-black iris.

### Detection of avian influenza antibodies in pale-versus dark-iris individuals

Prior the HPAI outbreak, all samples collected on healthy-like gannets in 2019 were seronegative for anti-NP and anti-H5 AIV antibodies (**Figure 2**). When exploring factors associated with the seropositive status of individuals determined using the H5-cELISA, the best model included year, 2 iris morphs and sex (**Supplementary Table 3**). The models considering three categories of iris specification were less parsimonious based on their AIC. In 2023, 100% (16/16) of the sampled individuals with a dark-iris phenotype had high levels of anti-NP AIV antibodies, among which 87.5% (14/16) were seropositive to H5 AIV (**Figure 2 and Supplementary Table 4**). Conversely, only a small fraction of individuals recorded with pale irises had antibodies against AIV: 9.52% (2/21) of seropositive birds to both NP and H5 AIV in 2023. In 2024, two years after the HPAI outbreak, anti-NP and anti-H5 AIV antibody levels of the sampled birds with a dark-iris phenotype remained high: 100% (13/13) and 69.2% (9/13) were seropositive to NP and H5 AIV respectively. Moreover, a fraction of sampled individuals with pale irises in 2024 had antibodies against AIV: 33.3% (5/15) and 26.7% (4/15) of seropositive birds to NP and H5 AIV respectively. The proportions did not differ between the sexes.

**Figure 2.**
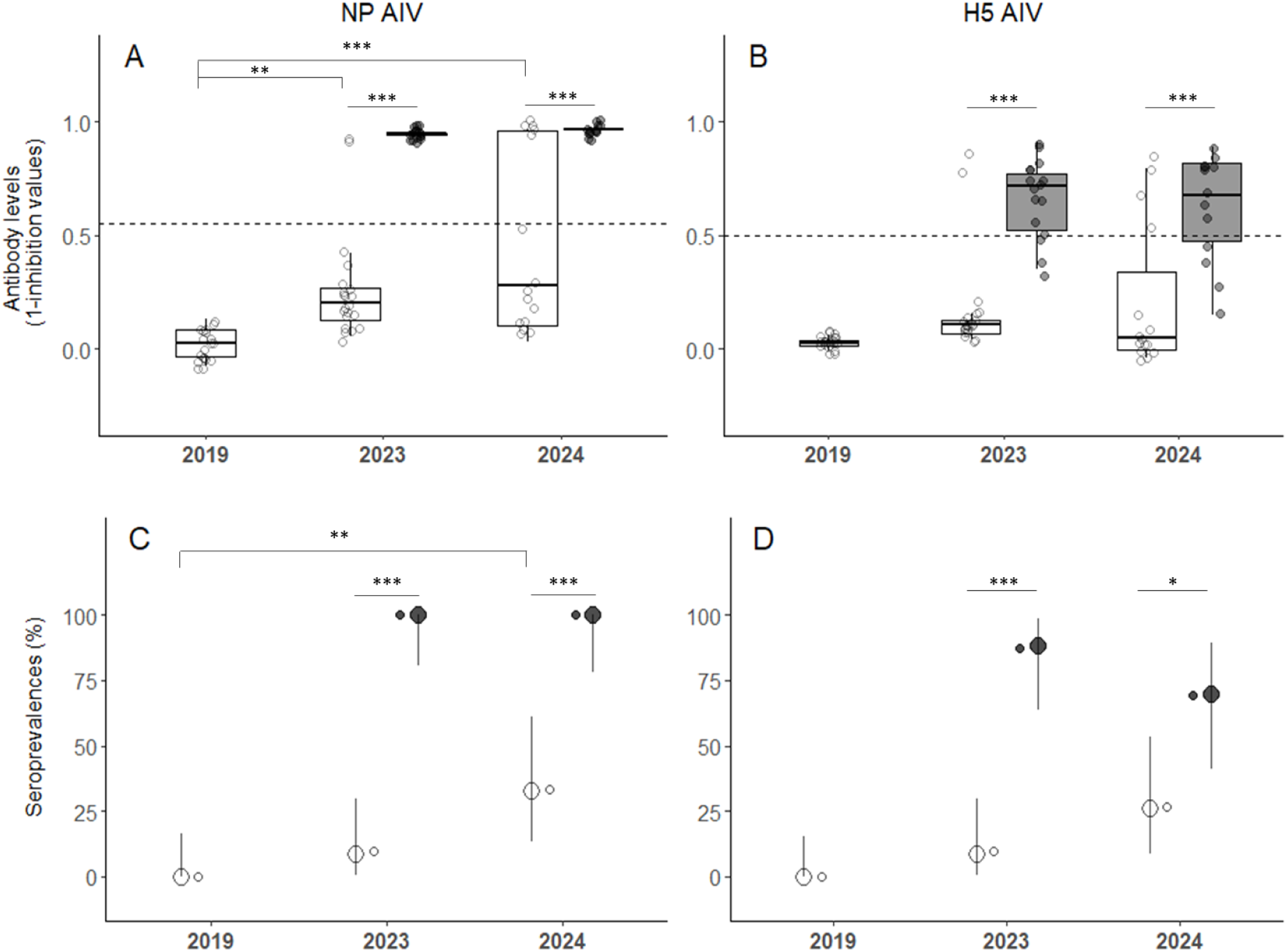
AIV antibody levels and associated seroprevalences measured in gannets sampled in 2019 (n = 20 birds), 2023 (n = 37 birds) and 2024 (n = 28 birds). Anti-NP (**A**) and anti-H5 (**B**) AIV antibody levels measured using competitive ELISAs and expressed as inhibition values. Black dashed lines refer to the inhibition values used as seropositivity thresholds. AIV-NP (**C**) and AIV-H5 (**D**) seroprevalences: circles and errors bars represent the true prevalence associated with the Clopper-Pearson 95% confidence intervals, while small dots refer to the observed prevalence. For all captions, healthy-like gannets with pale irises are represented using white, while the use of black refer to sampled gannets with a dark iris phenotype (with spotted-or full-black irises). Asterisks indicate the p-values resulting from the Two-Way analysis of variance tests (***, p < 0.001 ; **, p < 0.01 ; *, p < 0.05).

The results using the H5-iELISA were very similar, but a gradual association between the antibody status and the 3 iris phenotypes was detected (**Supplementary Table 3**). Full-black iris individuals showed higher antibody levels compared to spotted-black iris individuals, which themselves shoved higher antibody levels compared to pale iris individuals (**Supplementary Figure 3**).

All the juveniles sampled in the colony in 2023 were seronegative to both NP and H5 AIV (**Supplementary Figure 1**). Interestingly, among the 5 wrecked gannets with full-dark irises handled by the wildlife rescue center of Ile Grande in November 2023, only 3 birds were seropositive to H5 AIV (**Supplementary Figure 4**).

### Temporal persistence of antibody levels

Among the 11 individuals sampled in 2023 and recaptured in 2024, the correlation in their anti-H5 AIV antibody levels bet,ween the two years was high, with a slight decrease in 2024, but no change in serology status (**Figure 3**; Pearson correlation: r = 0.98, p < 0.001, anti-H5 antibody level (year+1) = 0.98189 * anti-H5 antibody level (year) - 0.06901). This suggests a high temporal persistence of antibody levels. Serological results obtained with the H,5 indirect ELISA (H5-iELISA) confirmed the high temporal persistence of anti-H5 antibody level in gannets (**Supplementary Figure 3C;** Pearson correlation: r = 0.91, p <0.001, anti-H5 antibody level (year+1) = 0.85956 * anti-H5 antibody level (year) + 0.05507), in line with the strong correlation of anti-H5 antibody titres obtained with the indirect and competitive ELISA assays (**Supplementary Figure 2**; Pearson correlation between results from the H5-cELISA and H5-iELISA: r = 0.78, p < 0.001, n = 84 samples). Importantly, none of the seronegative individuals in 2023 became seropositive in 2024 (**Figure 3**).

**Figure 3.**
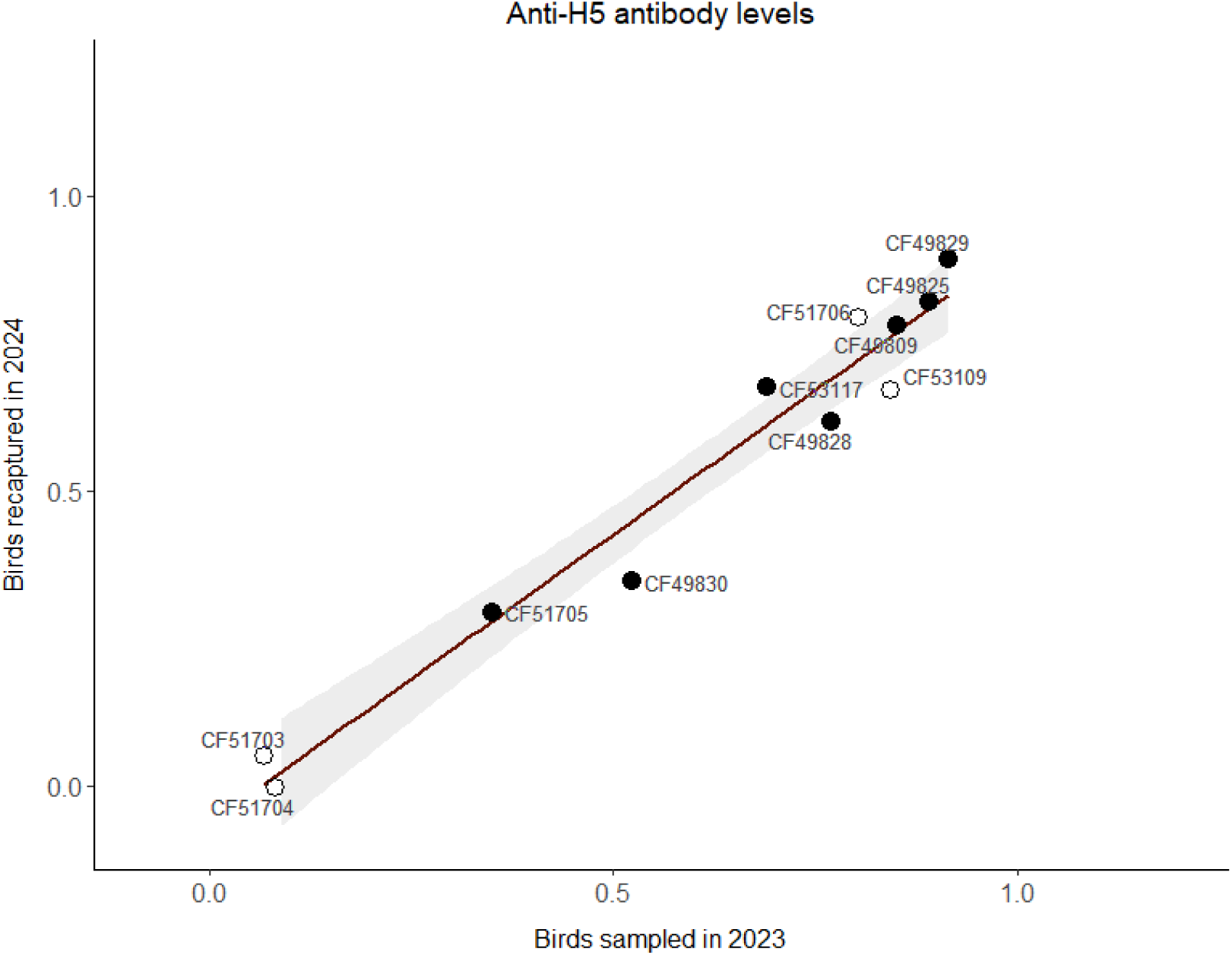
Anti-H5 AIV antibody levels measured in gannets sampled in 2023 and recaptured during the 2024 field session (n = 11 birds), using the H5 competitive ELISA assay (the plot displays 1-inhibition values). Healthy-like gannets with pale irises are represented with white circles (n = 4 birds), while black circles refer to gannets with a dark iris phenotype (spotted-or full-black irises, n = 7 birds). Each circle is associated with a unique alphanumeric code (e.g., CF49830), used to identify each sampled gannet. The red regression highlights a linear relationship between the anti-H5 AIV antibody level in gannets measured in 2024 as a function of that in 2003, associated to the following equation: anti-H5 antibody level (year + 1) = 0.98189 * H5 antibody level (year) – 0.06901. Pearson correlation r = 0.98, p < 0.001, n = 11 samples.

### HP H5 seroneutralisation

Seroneutralisation analyses conducted on plasma samples from 2023 and 2024 demonstrated that the capacity to neutralise the HP H5 virus was more efficient in gannets with a dark-iris phenotype compared to that of healthy-like gannets with pale irises (**Figure 4**, p < 0.01 and p < 0.001 respectively for the 2023 and 2024 season). Consistent titres were observed between the two years, with a slight decrease in 2024. Moreover, seroneutralisation titres were positively correlated with anti-H5 antibody levels, measured using either the H5-cELISA (**Figure 5**; Pearson correlation r = 0.80, p < 0.001, n = 51 samples), or the H5-iELISA (**Supplementary Figure 3D;** Pearson correlation r = 0.81, p < 0.001, n = 51 samples).

**Figure 4.**
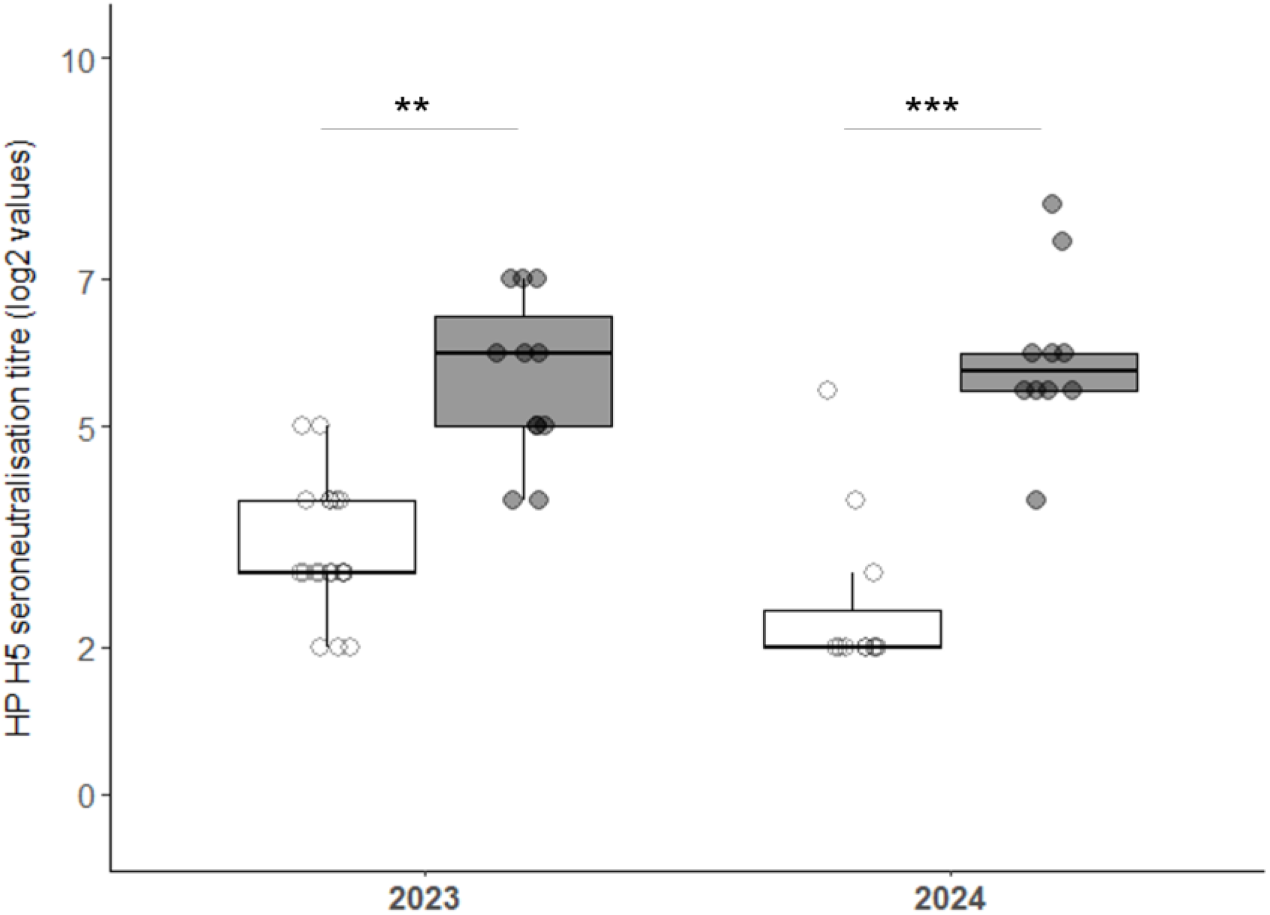
HP H5 seroneutralisation titres measured in gannet plasma samples collected in 2023 (n = 30 birds) and 2024 (n = 21 birds). White circles and boxes refer to gannets with healthy-like pale irises, while black circles and boxes refer to gannets with a dark iris phenotype (spotted-or full-black irises).

**Figure 5.**
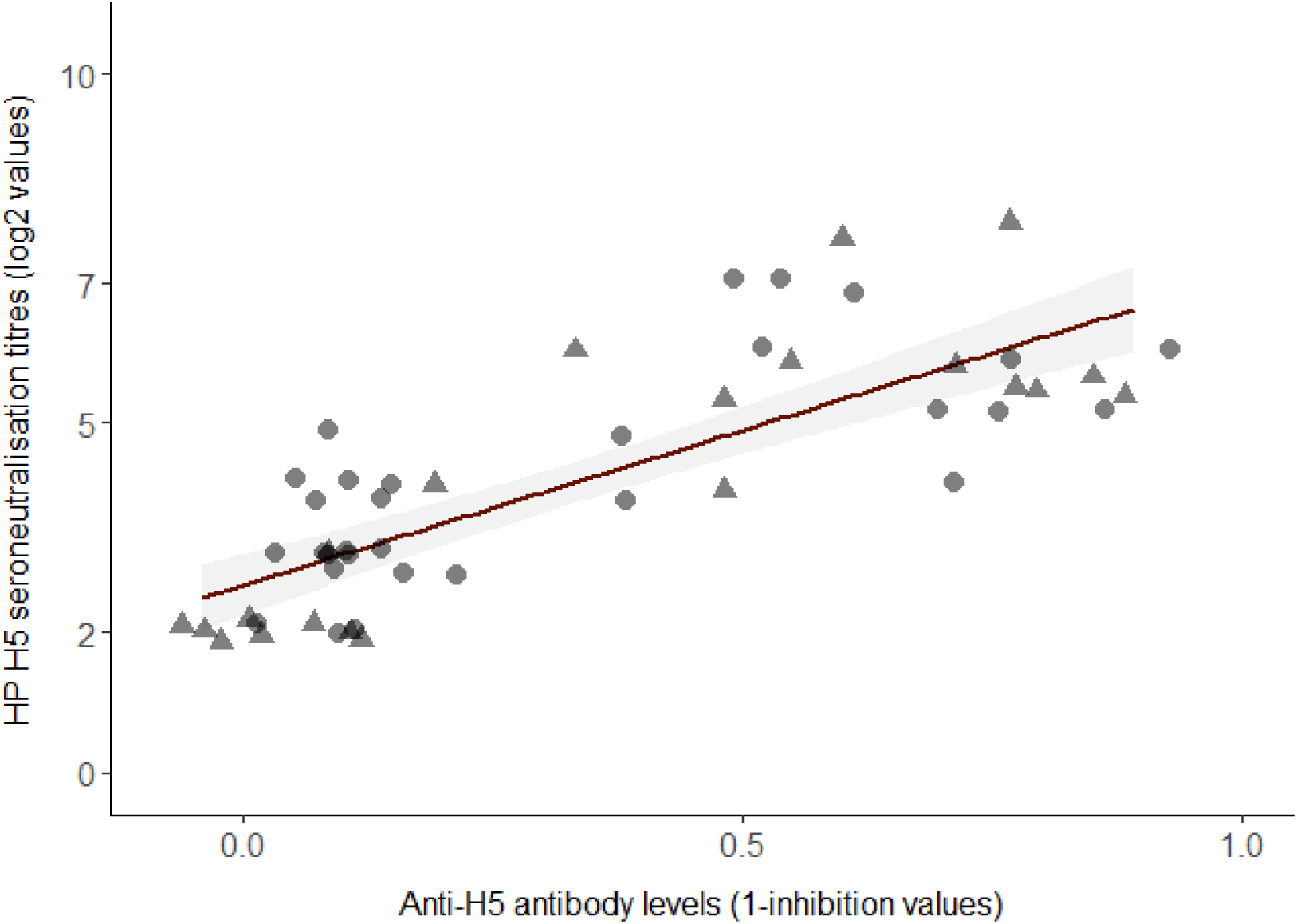
Correlation between HP H5 seroneutralisation titres and anti-H5 AIV antibody levels measured in gannet plasma samples using the H5 competitive ELISA assay. Circle dots indicate birds sampled in 2023 (n = 30 birds), while triangles refer to gannets sampled during the 2024 session (n = 21 birds). Pearson correlation r = 0.80, p < 0.001, n = 51 samples.

### Seroprevalences from proportions of dark iris individuals

Overall, based on H5 and NP seroprevalences in each group, and given that 22.3% of the breeding individuals were estimated to have a dark-iris phenotype in 2023, our results suggest that the overall proportion of seropositive individuals against H5 and NP antibodies in 2023 was circa 30% (% H5 seropositive in 2023 = 87.5% of the 22.30% dark-iris individuals + 9.52% of the 77.7% pale iris individuals = 19.51% + 7.40% = 26.91% H5 seropositive overall; % NP seropositive in 2023 = 100% of the 22.30% dark-iris individuals + 9.52% of the 77.7% pale iris individuals = 22.3% + 7.39% = 29.69% NP seropositive overall). Similarly in 2024, the overall proportions of seropositive birds were estimated to be 37% and 49% for H5 and NP antibodies respectively (% H5 seropositive in 2024 = 69.23% of the 23.7% dark-iris individuals + 26.67% of the 76.3% pale iris individuals = 16.40% + 20.34% = 36.74% H5 seropositive overall; % NP seropositive in 2024 = 100% of the 23.7% dark-iris individuals + 33.33% of the 76.3% pale iris individuals = 23.7% + 25.43% = 49.13% NP seropositive overall).

### Eco-epidemiological implications of the temporal persistence of immune resistance in interaction with annual survival

The model explorations showed that repeated outbreaks of HPAI lead to dramatic reduction of the population over a 20-year time span. This is especially the case if the annual adult survival rate is low (0.85), but also if the survival rate is high and the temporal persistence of immunity (OMEGA) is low (Figure 6 A and B). When 60% of the susceptible individuals were assumed to be infected during a year of HPAI outbreak (BETA = 0.6) and 30% were assumed to become immune after infection (GAMMA = 0.3), very low numbers of breeders were expected to remain after 20 years of repeated outbreaks if the proportion of individuals that remained immune between years was low (OMEGA = 0.1) compared to a scenario where this proportion is high (OMEGA = 0.99), as observed in our field data (**Figure 6A** versus **6B** in case of outbreaks every 4 years). If the annual survival of adults is high and the proportion of surviving adults that remain immune between years is high, the number of immune individuals remains relatively stable, and increases in proportion to the total population size (brown curve, **Figure 6A**). If the annual adult survival is low, as reported for the population of Rouzic before the HPAI outbreak (Grémillet et al. 2020), whatever the proportion of individuals that remain immune, a dramatic decrease in the population is predicted, leading fast to its extinction (**Figure 6C and D**). If the proportion of infected individuals that become immune resistant is low (low GAMMA) then the gannet population size drops to low level, regardless the proportion of individuals remaining immune resistant between 2 years (OMEGA)(**Supplementary Figure 5**). Conversely, when the proportion of infected individuals that becomes immune is high upon infection (high GAMMA), then the proportion of individuals remaining immune resistant between 2 years (OMEGA) has a strong positive effect on the population size (**Supplementary Figure 5**). The annual frequency of epizootics has little effect on the effect of the proportion of individuals remaining resistant on change in population size (**Supplementary Figure 6**). Overall, the model highlights the importance of the temporal persistence of immunity for populations to persist in the long-term after sporadic or regular epizootics events.

**Figure 6.**
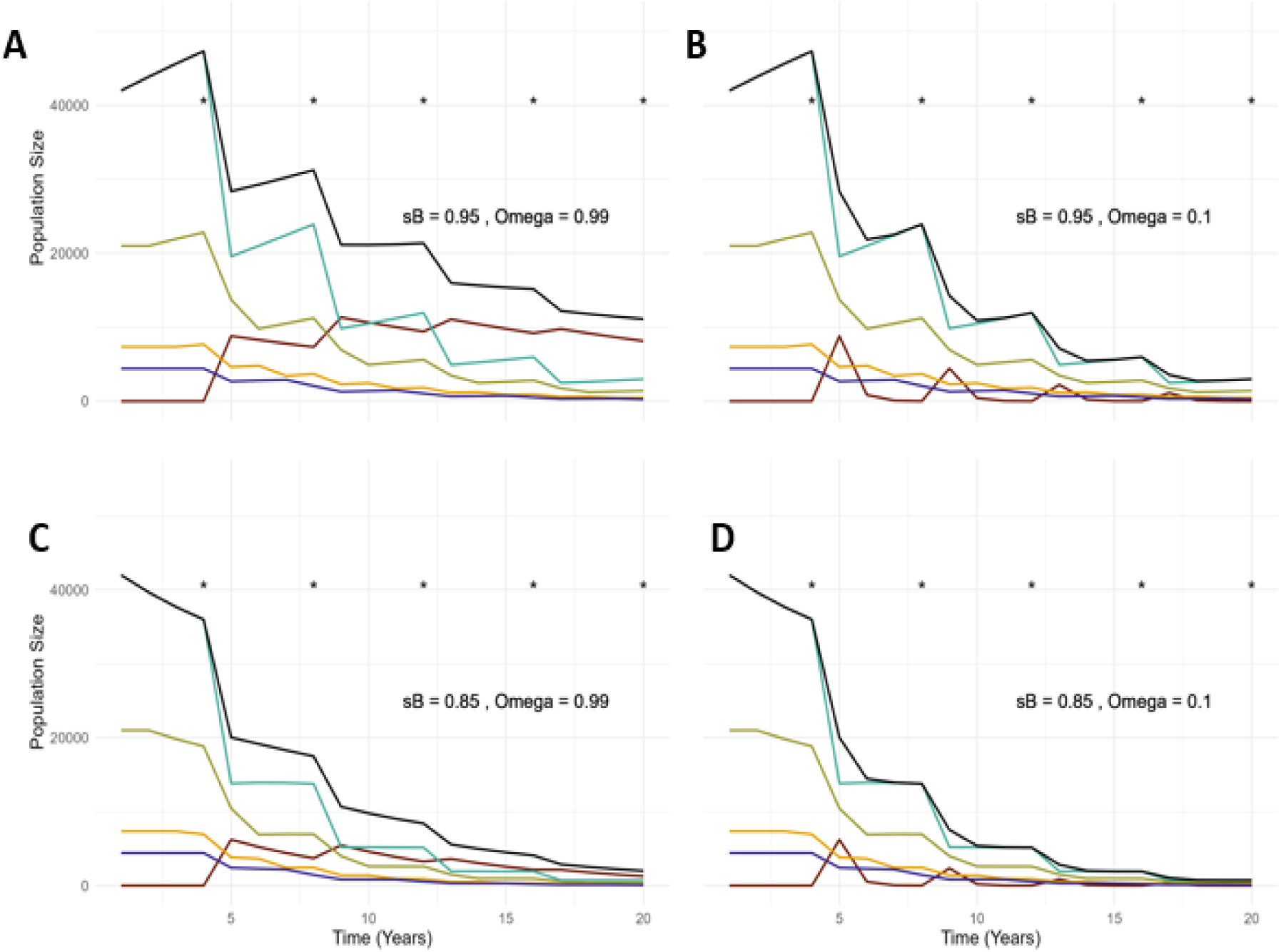
Change in the abundance of each demographic and epidemiological class of individuals over 20 years for populations affected every 4 years by an outbreak (*) for two levels of proportion of individuals that remain immune between years (OMEGA = 0.1 in (**A**) and (**C**) *versus* 0.99 in (**B**) and (**D**)) and adult survival (sB = 0.95 in (**B**) and (**D**) *versus* 0.8 in (**A**) and (**C**)); other parameters were fixed with BETA = 0.6 and GAMMA = 0.3. Colour lines show abundance of the total population (black), susceptible breeding individuals (light blue), 1-year-old immatures (beige), 2-year-old immatures (yellow), 3-year-old immatures (dark blue) and resistant (immune) adults (brown).

## Discussion

### High inter-year persistence of antibody levels

A very high proportion of individuals with antibodies against H5 avian influenza protein was detected among breeders with a dark-iris phenotype (full-or spotted-black iris) one year after the outbreak on Rouzic colony. Only a low proportion of breeding individuals had black eyes on the colony (circa 20%), and among those with pale-iris tested, a low proportion had anti-H5 antibodies. Levels of anti-H5 AIV antibody decreased very little between 2023 and 2024, with no loss of seropositive status among the 11 resampled individuals. Those results confirmed that the dark-iris phenotypic condition is associated with a history of past infection by HPAI virus in gannets, as suggested by Lane et al. (2023). Most importantly, they also show that anti-H5 antibodies highly persisted between years in that species. Further, we showed that the ability to seroneutralise a HPAI H5N1 2.2.3.4b lineage virus was also persistent and correlated, which suggests that the persistence of anti-H5 antibodies could be associated with the persistence of immune protection.

This important result is consistent with predictions that long-lived species should benefit from the evolution of long-lasting immune response (Garnier et al. 2013). From an epidemiological perspective, this has major implications for this long-lived species because it suggests that some segments of the population could be protected against the virus over the long term and that this proportion could increase after each outbreak. This could influence both the demographic consequences of future outbreaks and their probability of occurrence. From an evolutionary ecology perspective, these findings indicates that in a long-lived species such as the gannet, individuals could be selected to be protected against repeated exposure to the virus. It means also that the immunological landscape on which the virus evolves may strongly depend on the exposure history of individuals (Jamaleddine et al. 2026), which may be highly structured in space and time in such site-faithful species (Boulinier 2023).

### Eco-epidemiological implications of the temporal persistence of immunity

As the modelling approach highlighted, the persistence of immunity acquired in adults following exposure to the virus could mitigate the negative demographic impacts of subsequent outbreaks. However, the extent of immune protection is constrained by the annual survival rate of breeders, which ultimately determines how long immune individuals remain in the populations, and the proportion of individuals that become immune after infection. In a species with high annual adult survival, antibodies could still be detected in some individuals several years after an outbreak, but sustained exposure to HPAI could lead to rapid extinction of populations over a broad set of parameters, notably low annual survival rates. As for other long-lived species sensitive to HPAI, such as southern elephant seals or black-legged kittiwakes, the fact that the virus primarily affects adults leads to dramatic demographic effects in the short term (Bamford et al. 2025, Rømo et al. 2026), but also induce long recovery time for populations (Campagna et al. 2025, Lane et al. 2026).

Little data is available on antibody level persistence in wild birds following exposure to naturally occurring infectious agents (Staszewski et al. 2007, Gamble et al. 2019), notably in the case of avian influenza (Hill et al. 2016, 2019, Caliendo et al. 2022, White et al. 2024, Günther et al. 2024). Recent results in Indian Ocean seabirds suggest some but low inter-year consistency in serological status against avian influenza (antibody against the NP AI protein) between years (Lebarbenchon et al. 2023). The detection of antibodies has been suggested to be used as a surveillance tool (e.g., Kistler et al. 2012, 2014, Niqueux et al. 2014), but information about temporal persistence of antibody level is needed for interpreting results. In wild species, serological monitoring data is often cross sectional, with no repeated sampling of individuals (Wille et al. 2023, Greco et al. 2025, Knief et al. 2026, Rahman et al. 2026), which limits strongly the inference that can be made (Gamble et al. 2020). Under various assumptions, antibody levels can be used to infer epidemiological dynamics using model fitting to field data, but there too knowledge about antibody dynamics following exposure is paramount (Pepin et al. 2017, Wilber et al. 2017). The temporal persistence is expected to vary strongly among species, having been reported as low in ducks (White et al. 2024), but potentially long in flamingos following vaccination against HP H5 virus using an inactivated vaccine (Fernandez-Bellon et al. 2017). Recent results showing persistent antibody levels against H5 AIV protein in king penguin chicks more than 250 days after vaccination with an mRNA vaccine also suggest that long temporal persistence of antibody levels may occur in seabird species (Lejeune et al. 2026). In our case, the fact that low levels of antibodies were maintained in 2024 in all individuals that had low levels in 2023, suggests that there was no novel circulation of HPAI viruses on the colony, hence that the antibody levels measured in 2024 are likely to result from their initial exposure in 2022.

The fact that all juveniles were seronegative suggests that they had not been exposed to avian influenza virus since hatching, which, together with a lack of detection of abnormal mortalities, also represent evidence that there was no circulation of HPAI virus on the colony in 2023. After one month of age, nestlings would not be expected to retain maternal antibodies acquired from their mother through the egg yolk if their mother was seropositive (Garnier et al. 2012). Although the protection of young chicks through the transfer of anti-avian influenza maternal antibodies is unclear (e.g., Mass et al. 2011), the long-term persistence of antibody levels in adult gannet could have implications for the protection of young chicks after an outbreak (Garnier et al. 2012, Ramos et al. 2014). It is interesting to note that McLaughlin et al. (2025) reported proportions of gannet eggs containing antibodies against AIV NP (40%) and H5 proteins (20%) in 2023 on the colony of Bonaventure Island, Canada, which was greatly affected by a HPAI outbreak in 2022. No anti-AIV antibodies were detected in eggs before 2022 on that colony while eggs from several nests with dark iris attending adults had anti-influenza antibodies after the outbreak.

In this study, we have not considered factors that could affect the transmission rate of the virus among individuals or limit the occurrence of new outbreaks, in particular in relation to the proportion of individuals becoming immune, and thus to potential herd immunity effects (Keeling & Rohani 2012). Herd immunity could limit the occurrence of novel outbreaks in case of virus re-introduction, which should depend on the proportions of immune individuals. A critical knowledge gap is nevertheless whether the transmission of the virus is mostly occurring at sea or on the colony, and whether successful transmission is frequency or density dependent. Our simple model allowed us to highlight that in case of long temporal persistence of immunity, the proportion of immune individuals among breeding adults could increase relatively fast the proportion of immune individuals in case of repeated outbreaks, which could limit negative demographic impacts. With respect to the potentially protective effect of high levels of antibodies, it was interesting to detect intermediate levels of antibodies against H5 AIV in gannets with black irises that had been brought to the wildlife rescue center, potentially explaining their bad health condition. Several eco-epidemiological factors also need to be considered to fully interpret the implications of such a result, notably the complexity of immune responses in relation to the dynamics of exposure to a diversity of virus strains (Latorre-Margalef et al. 2013, Pepin et al. 2017).

### Implications for conservation

Although individuals with dark irises have a high probability of having avian influenza antibodies, several individuals with healthy-like iris phenotypes were seropositive to H5. Therefore, the presence of dark iris gannets is indicative of past exposure and survival to HPAI, but should be used with care for disease surveillance and population monitoring of gannets (Lane et al. 2023, Petalas et al. 2025). Our results have implications for the future dynamics of the French population, and more broadly, for other gannet populations that were massively affected by HPAI in 2022 (Grémillet et al. 2020, Jeglinksi et al. 2023, Matthiopoulos et al. 2026). They show that beyond causing substantial mortality among breeding adults, the HPAI outbreak of 2022 dramatically reshaped the composition of the population. It likely generated a significant proportion of newly immune individuals that could be temporally maintained in the population. This could directly impact the future of populations under pressure from HPAI, although the role of transmission of avian influenza viruses from other host species (Matthiopoulos et al. 2026) and at sea (Boulinier 2023), and possible viral antigenic drift, would have to be considered in attempts to propose predictions.

## Conclusion

Overall, the results suggest that naïve breeding populations of gannets exposed to the HPAI virus suffered high losses, but that after a single outbreak about 30-40 % of individuals developed humoral immunity against the virus that lasted at least two years, and that surviving breeding individuals that developed a dark-iris phenotype had acquired humoral immunity. Despite the mounting of a persisting and potentially protective immune response in some exposed individuals that survived, HP AI represents a critical threat for seabirds like gannets, already under pressure from other aspects of global change (Dias et al. 2019). Exploring the persistence of antibody levels in other long-lived species with various life-history traits would be important to confirm whether this is a broadly shared trait.

## Acknowledgements

We thank Nicolas Courbin, Tobie Getti, Grégoire Delavaud and Inès Mercereau for help at various stages of the study. Martha MacCall is thanked for conducting the molecular sexing, which was implemented at plateforme GEMEX. Funding support was provided by Ailes Marines, Ceva Wildlife Research Fund, CNRS Ecology Evolution SEE-Life program for long term monitoring, SO ECOPOP of OSU OREME, and ANR projects ECOPATHS (ANR-21-CE35-0016) and WILDFLU (ANR-25-CE35-0691). The study also benefited from the parallel implementation of French Polar Institute project IPEV ECOPATH-1151.

